# Modularity of biological systems: a link between structure and function

**DOI:** 10.1101/2023.09.11.557227

**Authors:** Claus Kadelka, Matthew Wheeler, Alan Veliz-Cuba, David Murrugarra, Reinhard Laubenbacher

## Abstract

This paper addresses two topics in systems biology, the hypothesis that biological systems are modular and the problem of relating structure and function of biological systems. The focus here is on gene regulatory networks, represented by Boolean network models, a commonly used tool. Most of the research on gene regulatory network modularity has focused on network structure, typically represented through either directed or undirected graphs. But since gene regulation is a highly dynamic process as it determines the function of cells over time, it is natural to consider functional modularity as well. One of the main results is that the structural decomposition of a network into modules induces an analogous decomposition of the dynamic structure, exhibiting a strong relationship between network structure and function. An extensive simulation study provides evidence for the hypothesis that modularity might have evolved to increase phenotypic complexity while maintaining maximal dynamic robustness to external perturbations.

## Introduction

Building complicated structures from simpler building blocks is a widely observed principle in both natural and engineered systems. In molecular systems biology, it is also widely accepted, even though there has not emerged a clear definition of what constitutes a simple building block, or module. Consequently, it is not clear how the modular structure of a system can be identified, why it is advantageous to an organism to be composed of modular components, and how we could take advantage of modularity to advance our understanding of molecular systems (*1–3*). In the (graph-theoretic) network representation of molecular systems, such as gene regulatory networks or protein-protein interaction networks, a module is typically considered to be a “highly” connected region of the graph that is “sparsely” connected to the rest of the graph, otherwise known as a community in the graph. Graph theoretic algorithms that depend on the choice of parameters and the specific definition of “highly” and “sparsely” are typically used to define modules (*4, 5*). Similar approaches are used for identifying modules in co-expression networks based on clustering of transcriptomics data (*6*).

A major limitation of this approach to modularity is that it focuses entirely on a static representation of gene regulatory networks and other systems. However, living organisms are dynamic, and need to be modeled and understood as dynamical systems. Thus, modularity should have an instantiation as a dynamic feature, as advocated in (*7*). The most common types of models employed for this purpose are systems of ordinary differential equations and discrete models such as Boolean networks and their generalizations, providing the basis for a study of dynamic modularity. In recent years, there have been an increasing number of papers that take this point of view. The authors of (*8*) argue that dynamic modularity may be independent of structural modularity, and they identify examples of multifunctional circuits in gene regulatory networks that they consider dynamically modular but without any underlying structural modularity. A similar argument is made in (*9*) by analyzing a small gene regulatory network example. For another example of a similar approach see (*10*).

The literature on how modularity might have evolved and why it might be useful as an organizational principle cites as the most common reasons robustness, the ability to rapidly respond to changing environmental conditions, and efficiency in the control of response to perturbations (*2, 11, 12*). An interesting hypothesis has been put forward in (*3*), namely that a modular organization of biological structure can be viewed as a symmetry-breaking phase transition, with modularity as the order parameter.

This literature makes clear that research on the topic of modularity in molecular systems, both structural and dynamic, would be greatly advanced by clear definitions of the concept of module, both structural and dynamic. This would in particular help to decide whether and how structural and dynamic modularity are related, and it would provide a basis on which to distinguish between modularity and multistationarity of a dynamic regulatory network. To be of practical use, such a theory should include algorithms to decompose a dynamic network into structural and/or dynamic modules. At the same time, it would be of great practical value, for instance for synthetic biology, to understand how systems can be composed from modules that have specific dynamic properties.

The search for such algorithms has led us to look for guidance to mathematics, as a complement to biology. After all, if the dynamic mathematical models that are widely used to encode gene regulatory networks are appropriate representations, and if modularity is indeed an important feature of such networks, then it should be reflected in the model structure and dynamics. Choosing the widely-used modeling framework of Boolean networks, we asked whether it is possible to identify meaningful concepts of modularity that, ideally, link both the structural and dynamic aspects. Modularity is fundamentally about connectivity. The central dynamic instantiation of connectivity is the feedback loop, which we therefore choose as the defining feature. The concept of module we propose is structural, in terms of special subgraphs of the (directed) graph of dependencies of network nodes. These subgraphs, called strongly connected compoments (SCC), are maximal with respect to the property that every node is connected to every other node in the subgraph through a directed path. In other words, none of the nodes in the SCC are involved in feedback loops that are not entirely within the SCC.

The main result of this paper is that this structural decomposition of the model into modules induces a similar decomposition of model dynamics, explicitly linking the dynamics of the structural modules in a mathematically clearly specified way. This theorem links structural and dynamic modularity, and provides an example of how network structure influences network function. We provide an important application of this theorem to network control by showing that in order to control a network, it is sufficient to control its modules, and we provide an application of this result to a published cancer signaling network. This result is important both for applications to, e.g., medicine, and might provide a candidate for a mechanisms that allows organisms to quickly respond to changes in their external environment. We also discuss our results in the context of published Boolean network models of regulatory networks and provide specific instantiations of our decomposition theorem. Finally, we address the question as to why evolution should favor modularity as a structural and dynamic feature. We carry out an extensive simulation study that provides evidence for the hypothesis that modularity enables phenotypic complexity while maintaining maximal robustness to external perturbations.

### Boolean networks

For the purpose of this article, we will focus on the class of Boolean networks as a modeling paradigm. Recall that a Boolean network *F* on variables *x*_1_, …, *x*_*n*_ can be viewed as a function on binary strings of length *n*, which is described coordinate-wise by *n* Boolean update functions *f*_*i*_. Each function *f*_*i*_ uniquely determines a map

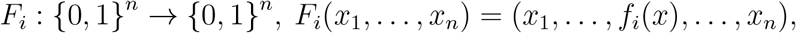

where *x* = (*x*_1_, …, *x*_*n*_). Every Boolean network defines a canonical map, where the functions are synchronously updated:

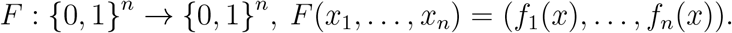

In this paper, we only consider this canonical map, i.e., we only consider synchronously updated Boolean network models. Two directed graphs can be associated to *F* (see Fig. 1 for an example). The *wiring diagram* (also known as dependency graph) contains *n* nodes corresponding to the *x*_*i*_, and has a directed edge from *x*_*i*_ to *x*_*j*_ if *f*_*j*_ depends on *x*_*i*_. The *state space* of *F* contains as nodes the 2^*n*^ binary strings, and has a directed edge from **u** to **v** if *F* (**u**) = **v**. Each connected component of the state space gives an *attractor basin* of *F*, which consists of a directed loop, *the attractor*, as well as trees feeding into the attractor. Attractors can be steady states (also known as fixed points) or limit cycles. Each attractor in a biological Boolean network model typically corresponds to a distinct phenotype (*13*). The *set of attractors* of *F*, denoted 𝒜(*F*), contains all attractors, i.e., all minimal subsets 𝒞 ⊆ {0, 1}^*n*^ satisfying *F* (𝒞) = 𝒞. Note that a limit cycle of length *k* represents *k* trajectories. For example, the 2-cycle (010, 101) in Fig. 1 represents (010, 101, 010, …) and (101, 010, 101, …). This distinction becomes important later, when decomposing the dynamics of Boolean networks.

**Figure 1:**
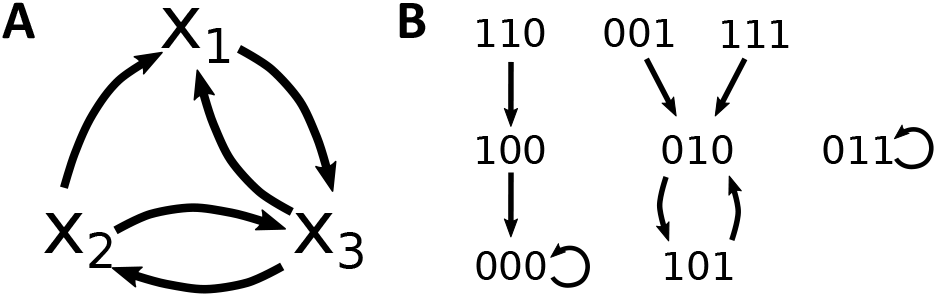
Wiring diagram and state space of the Boolean network *F* = (*f*_1_, *f*_2_, *f*_3_) = (*x*_2_ ∧ *¬x*_3_, *x*_3_, *¬x*_1_ ∧ *x*_2_). (a) The wiring diagram encodes the dependency between variables. Subnetworks are defined on the basis of the wiring diagram. For example, the subnetwork 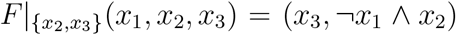 is the restriction of *F* to {*x*_2_, *x*_3_} and contains external parameter *x*_1_. (b) The state space is a directed graph with edges between all states and their images. This graph therefore encodes all possible trajectories and attractors. Here, *F* has two steady states, 000 and 011, and one limit cycle, (010, 101), so *A*(*F*) = {000, 011, (010, 101)}.

### A structural definition of modularity for Boolean networks

Given a Boolean network *F* and a subset *S* of its variables, we can define a subnetwork of *F*, denoted *F* |_*S*_, as the restriction of *F* to *S*. If some variables in *S* are regulated by variables not in *S*, then we require these regulations to be included in *F* |_*S*_. In this case, the subnetwork is a

Boolean network with external parameters. For the example in Fig. 1, the subnetwork 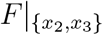 contains *x*_1_ as external parameter because *x*_1_ regulates *x*_3_. If the variables in *S* form a SCC (that is, (i) every pair of nodes in *S* (excluding possible external parameters) is connected by a directed path, and (ii) the inclusion of any additional node in *S* will break this property), we call the subnetwork a *module*.

The wiring diagram of any Boolean network *F* is either strongly connected or it consists of a collection of SCCs where connections between two SCC point in only one direction. Let *W*_1_, …, *W*_*m*_ be the SCCs of the wiring diagram, with *Y*_*i*_ denoting the set of variables in SCC *W*_*i*_ (note ∪_*i*_*Y*_*i*_ = *Y* and *Y*_*i*_ ≠ *Y*_*j*_ for *i* ≠ *j*). Then, the *modules* of *F* are 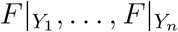, the restrictions of *F* to the *Y*_*i*_. By setting *W*_*i*_ → *W*_*j*_ if there exists at least one edge from a node in *W*_*i*_ to a node in *W*_*j*_, we obtain a directed acyclic graph

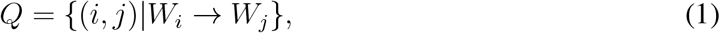

which describes the connections between the modules of *F* .

As we will show later, any Boolean network can be decomposed into modules and this structural decomposition implies a decomposition of the network dynamics, which is of practical utility. The main question to be answered at this point, though, is whether there exists biological evidence that our concept of modularity and the structural and dynamic decomposition theory that follows does in fact reflect reality.

### Modularity in expert-curated biological networks

A recent study investigated the features of 122 distinct published, expert-curated Boolean network models (*14*). Analyzing the wiring diagrams of these models, we found that almost all of them (113, 92.6%) contained at least one feedback loop and thus at least one non-trivial SCC/module (which contains more than one node). The nine models that only contained single-node SCCs mainly describe signaling pathways. Thirty models (24.6%) contained even more than one non-trivial SCC, with one Influenza A virus replication model possessing eleven (*15*). The directed acyclic graph structure (Eq. 1) of these models varied widely (Fig. S1). While the average connectivity of a network was not correlated with the number of non-trivial SCCs (*ρ*_Spearman_ = 0.05, *p* = 0.61), network size was positively correlated (*ρ*_Spearman_ = 0.35, *p <* 10^−4^). The same trends persisted when considering the binary variable “multiple non-trivial SCCs” (multivariable logistic regression: connectivity *p* = 0.36, size *p* = 0.003).

Modules are subnetworks that carry out key control functions in a cell. It would therefore not be surprising if there was a selection bias among systems biologists to focus their attention on such modules. Larger networks are still challenging to build and analyze since an accurate formulation of a biological network model requires a substantial amount of data for a careful inference and calibration of the update rules by a subject expert (*16, 17*). For this reason most published expert-curated models might focus on one specific cellular function of interest and contain therefore only one non-trivial SCC.

### Modularity confers phenotypical robustness and a rich dynamic repertoire

To provide additional evidence that SCCs form biologically meaningful modules, we performed a computational study which shows that the presence of several modules confers robust phenotypes and a rich dynamic repertoire, both desirable features for an organism.

Biological networks must harbor multiple phenotypes, allowing the network to dynamically shift from one attractor to another based on its current needs. This shift is typically mitigated by external signals. Many evolutionary innovations are the result of newly evolved attractors of GRNs (*18,19*). The number of attractors of a Boolean network therefore describes its *dynamical complexity*.

Furthermore, biological networks need to robustly maintain a certain function (i.e., phenotype) in the presence of intrinsic and extrinsic perturbations (*20, 21*). At any moment, these perturbations may cause a small number of genes to randomly change their expression level. For a Boolean GRN model, this corresponds to an unexpressed gene being randomly expressed, or vice versa. The robustness of the network describes how a perturbation on average affects the network dynamics. One popular robustness measure for Boolean networks, the Derrida value, describes the average Hamming distance between two states after one synchronous update according to the Boolean network rules, given that the two states differed in a single node (*22*). Due to the finite size of the state space, any state of a BN eventually transitions to an attractor, which corresponds to a distinct biological phenotype. Thus, the Derrida value is a meaningful robustness measure

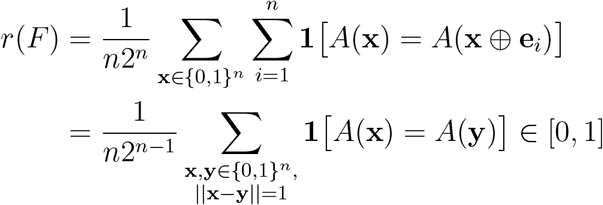

Here, **e**_*i*_ is the *i*th unit vector and *A*(**x**) labels the attractor that state **x** transitions to. Geometrically, if we consider the Boolean hypercube with each vertex in {0, 1}^*n*^ labeled by the attractor that the vertex-associated state eventually transitions to, then *r*(*F*) is the proportion of edges, which connect vertices with the same value.

Clearly, *r*(*F*) = 1 if a Boolean network *F* possesses only a single attractor. Moreover, the expected value, 𝔼 [*r*(*F*)], decreases as the number of attractors of *F* increases. This implies that the phenotypical robustness and the dynamical complexity are negatively correlated and that there exists a trade-off when trying to maximize both. It is reasonable to hypothesize that evolution favors robust GRNs that give rise to sufficient variety in the phenotype space. In line with this, we hypothesized that modular networks have higher robustness than non-modular networks with the same dynamical complexity.

To test this hypothesis, we generated nested canalizing Boolean networks with *N* = 60 nodes, a fixed in-degree of 3, and *m* = 1, …, 6 modules (i.e., SCCs of the wiring diagram) of size *N/m*. Networks with more modules possessed on average a higher dynamical complexity, quantified here as the number of attractors (Fig. 2A). At a fixed dynamical complexity, the more modular a network the higher was its average phenotypical robustness (Fig. 2B). This finding supports the hypothesis that a modular design serves as an evolutionary answer to a multi-objective optimization problem.

**Figure 2:**
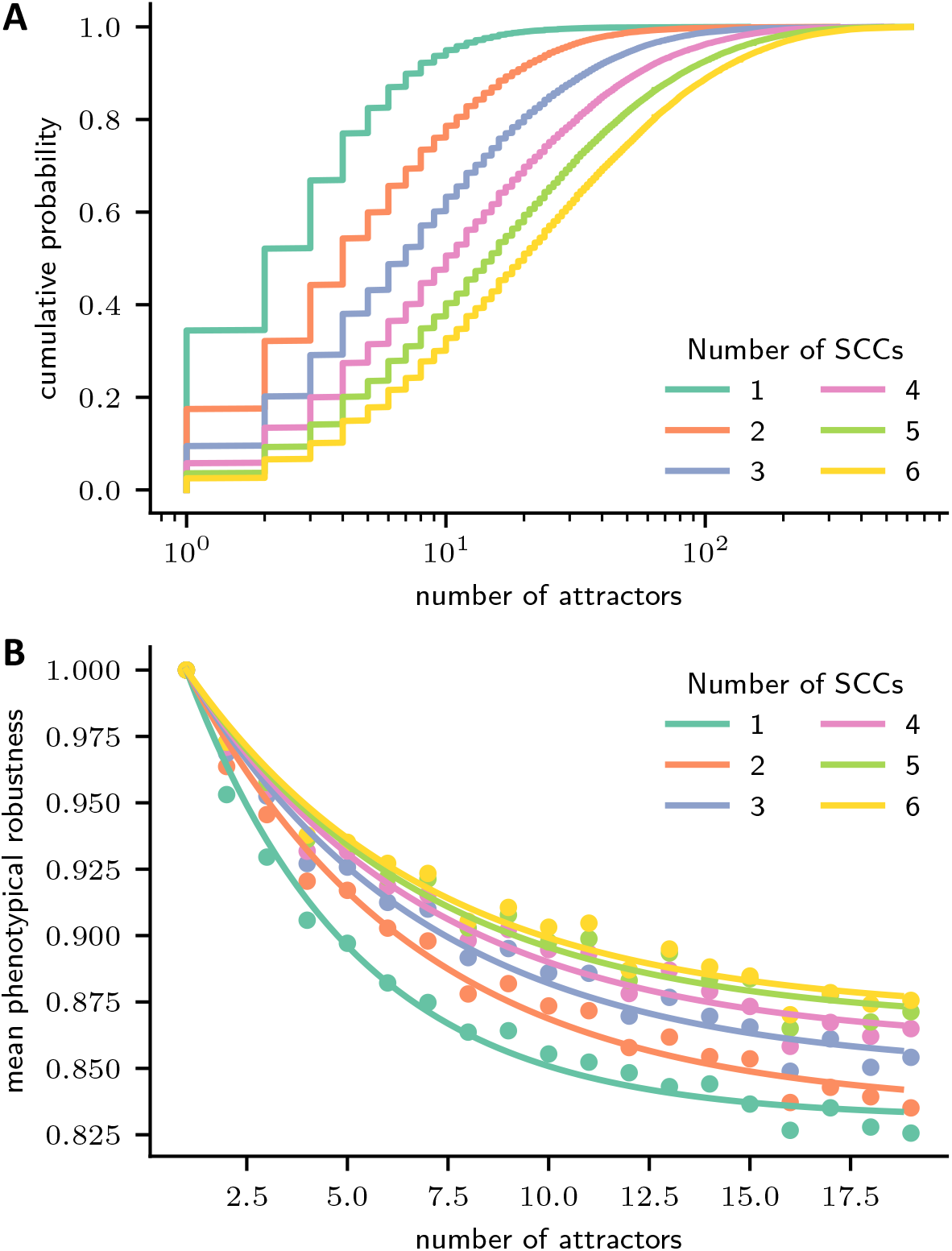
Modularity confers dynamical complexity and phenotypical robustness. 60-node nested canalizing Boolean networks with a constant in-degree of 3 and with 1-6 modules (i.e., SCCs of the wiring diagram) of equal size were generated (50,000 networks each). For each modular network, a weakly connected directed graph describing the connections between modules, as well as a single edge connecting an upstream with a downstream module were selected uniformly at random. By following the transitions of 500 random initial states to their attractors, the phenotypical robustness and a lower bound for the dynamical complexity (here, number of attractors) were established for each network. (A) Cumulative empirical density function of the number of attractors, stratified by the number of modules or SCCs. (B) The mean phenotypical robustness (*y*) is plotted against the number of discovered attractors (*x*), stratified by the number of modules or SCCs (dots). Since *y*(1) = 1, the two-parameter function *y* = *α*+(1−*α*)*e*^−*k*(*x*−1)^ is fitted to the means of the number of attractors for *x* = 1, …, 19 (lines).

### Structural decomposition of Boolean networks

Thus far, we have described how to define modules as restrictions of Boolean networks, and provided evidence that modules defined this way are biologically meaningful. To obtain a successful decomposition theory, we also require the inverse operation of a restriction: a semidirect product that combines two Boolean networks, *F* and *G*, such that *F* is the upstream module and *G* is the downstream module. The *coupling scheme P* contains the information which nodes in *F* regulate which nodes in *G*. We denote the combined Boolean network as *F* ⋊_*P*_ *G* and refer to this as the coupling of *F* and *G* by the coupling scheme *P* or as the semi-direct product of *F* and *G* via *P* (detailed definition i n SI Appendix, section 1). (The motivation for the term “semi-direct product” comes from the fact that the combination of the two subnetworks is like a product, except that *F* acts on *G* through *P*, which is not the case in an actual product. The term is also used in mathematical group theory, which provided the motivation for our decomposition approach.)

As an example, consider the Boolean networks *F* (*x*_1_, *x*_2_) = (*x*_2_, *x*_1_) and *G*(*u*_1_, *u*_2_, *y*_1_, *y*_2_) = (*u*_1_ ∨ (*u*_2_ ∧ *y*_2_), *¬u*_2_ ∧ *y*_1_) where *G* possesses two external parameters, *u*_1_ and *u*_2_. With the coupling scheme *P* = {*x*_1_ → *u*_1_, *x*_2_ → *u*_2_}, we obtain the combined nested canalizing network *F* ⋊_*P*_ *G* : {0, 1}^4^ → {0, 1}^4^,

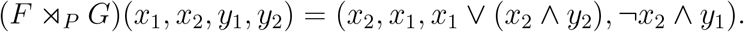

At the wiring diagram level, this product can be seen as the union of the two wiring diagrams and some added edges determined by the coupling scheme *P* (Fig. 3). If instead *G*(*u*_1_, *u*_2_, *y*_1_, *y*_2_) = *u*_1_ + *u*_2_ + *y*_2_, *u*_2_ + *y*_1_ with *F* and *P* as before, then we obtain the linear network

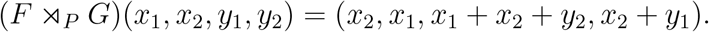

At the wiring diagram level, this product looks exactly the same (Fig. 3).

**Figure 3:**
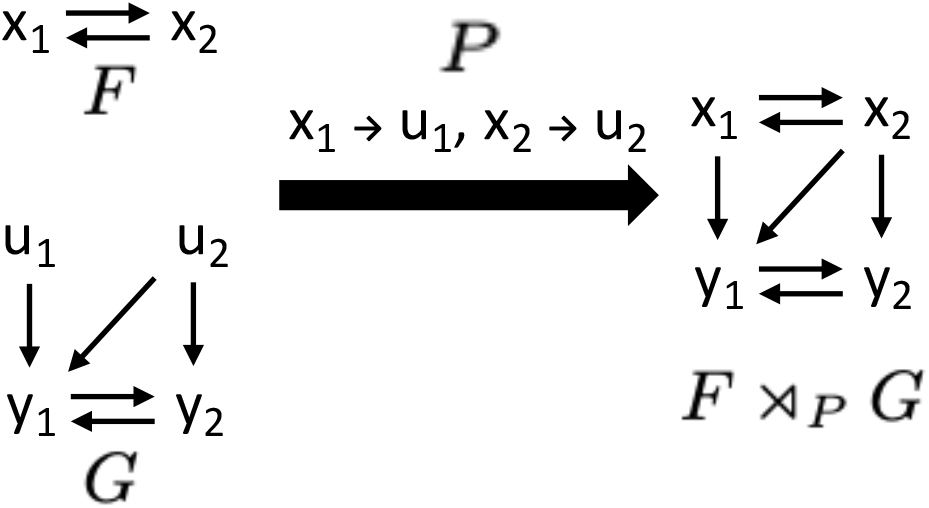
Semi-direct product of Boolean networks. Wiring diagrams of independent Boolean networks *F* and *G* (where *G* has external parameters) can be combined into *F* ⋊_*P*_ *G*, the semi-direct product of *F* and *G*. The coupling scheme *P* describes which variables of *F* take the place of the external parameters and act as inputs to *G*.

We can prove that every network is either a module or can be decomposed into a semi-direct product of two networks. That is, if a Boolean network *F* is not a module (i.e., if its wiring diagram is not strongly connected), then there exist *F*_1_, *F*_2_, *P* such that *F* = *F*_1_ ⋊_*P*_ *F*_2_, and we call such a network *F decomposable*. We can even find a decomposition such that *F* _1_ is a module. By induction on the downstream component *F*_2_, it follows that any Boolean network is either a module or decomposable into a unique series of semi-direct products of modules. That is, for any Boolean network *F*, there exist unique modules *F*_1_, …, *F*_*m*_ (*m* = 1 if *F* is itself a module) such that

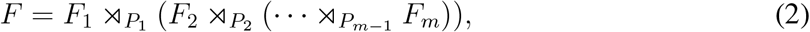

where this representation is unique up to a reordering, which respects the partial order induced by the directed acyclic graph *Q* (Eq. 1). The collection of coupling schemes *P*_1_, …, *P*_*m*−1_ depends on the particular choice of ordering, as well as on the placement of parentheses in the decomposition of *F*, which may be rearranged in any associative manner. SI Appendix, section 1 contains the proofs of these theorems.

### Dynamic decomposition of Boolean networks

When the variables of a network *F* can be partitioned such that *F* = *F*_1_ ⋊_*P*_ *F*_2_ = *F*_1_ *× F*_2_ is simply the cross product of two networks *F*_1_ and *F*_2_, i.e., the coupling scheme *P* = ∅, then the dynamics of *F* can be determined directly from the dynamics of *F*_1_ and *F*_2_. The dynamics of *F* consists of coordinate pairs (*x, y*) such that

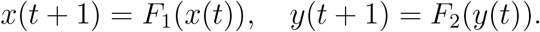

If trajectories 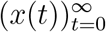 and 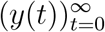 have periods *l* and *m*, respectively, then the periodicity of the trajectory 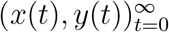 is the least common multiple of *l* and *m*. Moreover, the set of periodic points (i.e., attractors) of *F* is the Cartesian product of the set of periodic points of *F*_1_ and periodic points of *F*_2_.

For example, the Boolean network *F* (*x*_1_, *x*_2_, *x*_3_, *x*_4_) = (*x*_2_, *x*_1_, *x*_4_, *x*_3_) can be seen as *F* = *F*_1_ *× F*_2_, where *F*_1_(*x*_1_, *x*_2_) = (*x*_2_, *x*_1_) and *F*_2_(*x*_3_, *x*_4_) = (*x*_4_, *x*_3_). The sets of attractors of *F*_1_ and *F*_2_ are 𝒜 (*F*_1_) = {00, 11, (01, 10)} and 𝒜 (*F*_2_) = {00, 11, (01, 10)} (where we omit parentheses around steady states). By concatenating the attractors of *F*_1_ and *F*_2_, we obtain the attractors of *F* (Fig. 4A). Note that we have two ways of concatenating the limit cycle (01, 10) of *F*_1_ and the limit cycle (01, 10) of *F*_2_ to obtain attractors of *F* . In general, we have the following equation that formally states that attractors of *F*_1_ *× F*_2_ are given by concatenating attractors of *F*_1_ and *F*_2_.

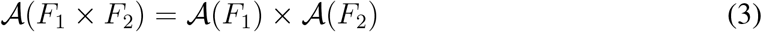

The computation of the attractors of *F* becomes more complicated when *F* is slightly modified so that *F* (*x*_1_, *x*_2_, *x*_3_, *x*_4_) = (*x*_2_, *x*_1_, *x*_2_*x*_4_, *x*_3_) = *F*_1_ ⋊_*P*_ *F*_2_, where *F*_1_ as before and *F*_2_ = (*ux*_4_, *x*_3_) with external parameter *u* and coupling scheme *P* = {*x*_2_ → *u*}. Since the coupling between *F*_1_ and *F*_2_ is no longer empty, not every combination of attractors of *F*_1_ and *F*_2_ will result in an attractor of *F* (Fig. 4B). For example, (01, 10) ∈ 𝒜 (*F*_1_) and (01, 10) ∈ 𝒜 (*F*_2_) do give rise to an attractor of *F*, while (01, 10) ∈ 𝒜 (*F*_1_) and (10, 01) ∈ 𝒜 (*F*_2_) do not. The set of attractors, 𝒜 (*F*), is the union of 00 *×* 00, 11 *× A*(*F*_2_), and (01, 10) *×* {00, (01, 10)}, and is thus a subset of the attractors of the Cartesian product (Fig. 4A). This is, however, not always the case but depends on the particular coupling between the networks. Hence, Eq. 3 is not valid in general.

**Figure 4:**
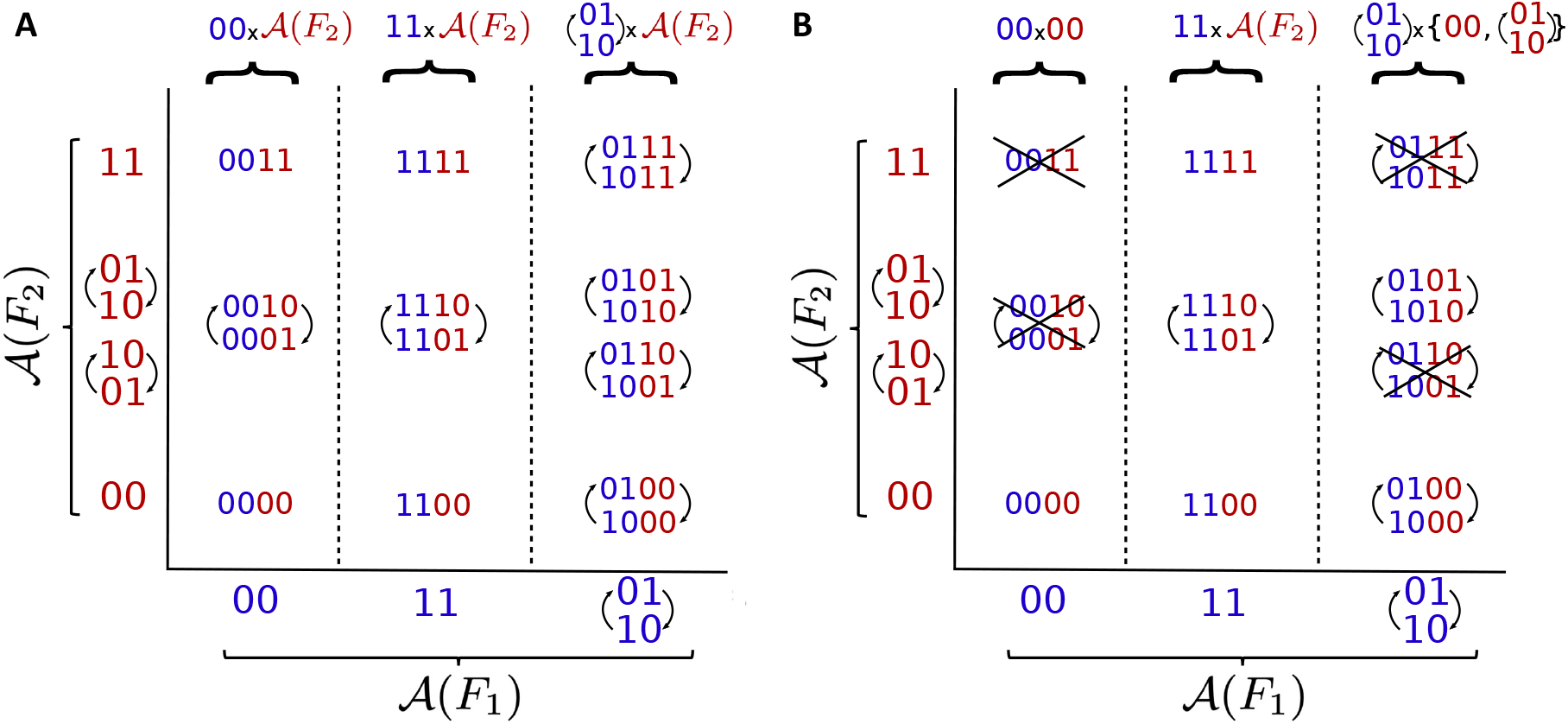
Attractors of a Cartesian product and a semi-direct product. (A) The space of attractors of a Cartesian product *F* = *F*_1_ *× F*_2_, with *F*_1_(*x*_1_, *x*_2_) = (*x*_2_, *x*_1_), *F*_2_(*x*_3_, *x*_4_) = (*x*_4_, *x*_3_), can be seen as a Cartesian product of 𝒜 (*F*_1_) and 𝒜 (*F*_2_). To illustrate the different ways to combine attractors of *F*_1_ and *F*_2_, in the panel we explicitly write (01, 10) and (10, 01) for *F*_2_. (B) In general, the coupling of networks does not behave as a Cartesian product and the space of attractors depends on this coupling. The crossed-out attractors indicate which attractors from the Cartesian product are lost when using a semi-direct product with coupling scheme *P* = {(*x*_3_, *x*_2_*x*_4_)}, and *F*_1_, *F*_2_ as in A.

In order to study the dynamics of decomposable networks, we need to understand how a trajectory, which describes the behavior of an “upstream” network at an attractor, influences the dynamics of a “downstream” network. The trajectory of an “upstream” *m*-node network *F*_1_ at an attractor 𝒞_1_ = (*α*_1_, …, *α*_*r*_) can be described by 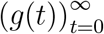, a sequence with elements in {0, 1}^*m*^. This trajectory has period *r*, the length of the attractor. The dynamics of the “downstream” *n*-node network *F*_2_ depend on *F*_1_. Therefore, *F*_2_ is a *non-autonomous Boolean network*,

defined by

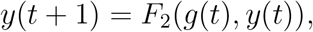

where *F*_2_ : {0, 1}^*m*+*n*^ → {0, 1}^*n*^. SI Appendix, section 2 contains a detailed definition and examples of non-autonomous Boolean networks. To make the dependence of *F*_2_ on the choice of upstream attractor *C*_1_ ∈ 𝒞_1_ explicit, we often write 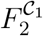 instead of simply *F*_2_. If 𝒞_2_ = (*β*_1_, …, *β*_*s*_) is an attractor of 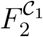, then

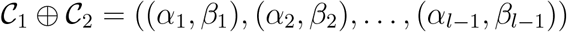

is an attractor of the combined network *F* = *F*_1_ ⋊_*P*_ *F*_2_ of length *l* := *lcm*(|𝒞_1_|, |𝒞_2_|), the least common multiple of |𝒞_1_| and |𝒞_2_|.

Iterating over all attractors of *F*_1_ (that is, all 𝒞_1_ ∈ 𝒜 (*F*_1_)) as well as all attractors of the corresponding non-autonomous networks 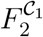 (that is, all 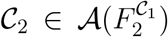 yields all attractors of the combined network *F* . After the structural decomposition theorem (Eq. 2), this dynamic decomposition theorem constitutes the second main theoretical result. Mathematically, it can be expressed as

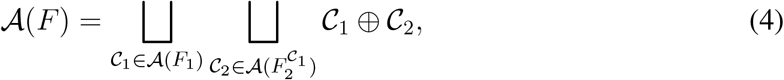

which can be written as 𝒜 (*F*_1_) ⋊_*P*_ 𝒜 (*F*_2_) to highlight the analogy between the structural decomposition of a Boolean network and the decomposition of its dynamics. With this, the dynamic decomposition theorem states 𝒜 (*F*_1_ ⋊_*P*_ *F*_2_) = 𝒜 (*F*_1_) ⋊_*P*_ 𝒜 (*F*_2_), which implies a distributive property for the dynamics of decomposable networks. Note that if *P* is empty, then 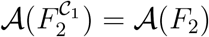 for all 𝒞_1_ and we recover Eq. 3, 𝒜 (*F*_1_ *× F*_2_) = 𝒜 (*F*_1_) *×* 𝒜 (*F*_2_).

The dynamics of a Boolean network *F*, which decomposes into modules *F*_1_, …, *F*_*m*_, can thus be computed from the dynamics of its modules. That is,

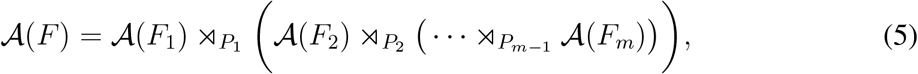

where the placement of the parentheses may be rearranged in any associative manner, just as for the structural decomposition in Eq. 2. SI Appendix, section 2 contains the proof of the dynamic decomposition theorem as well as instructional examples.

### Efficient control of decomposable Boolean networks

The state space of a Boolean network grows exponentially in the number of variables. Therefore, the decomposition theorems can reduce the time needed to perform various computations by orders of magnitude for networks with several larger modules. Besides an efficient strategy to compute all attractors of a Boolean network, the structural decomposition theorem can also be applied to efficiently identify controls of Boolean networks, a topic that has received recent attention (*23–25*). Drug developers wonder, for example, which nodes in a gene regulatory network need to be controlled by an external drug to ensure the network transitions to a desired phenotype, typically corresponding to a specific network attractor.

Two types of control actions are generally considered: edge controls and node controls. For each type of control, one can consider deletions or constant expressions, as defined in (*26*). The motivation for considering these control actions is that they represent the common interventions that can be implemented in practice. For instance, edge deletions can be achieved by the use of therapeutic drugs that target specific gene interactions, while node deletions represent the blocking of effects of products of genes associated to these nodes (*27, 28*).

A set of controls *μ stabilizes* a Boolean network at an attractor 𝒞 when the resulting network after applying *μ* possesses 𝒞 as its only attractor. As described in detail in (*29*), the decomposition into modules can be used to obtain controls for each module, which can then be combined to obtain a control for an entire network. Specifically, for a decomposable network *F* = *F*_1_ ⋊_*P*_ *F*_2_, if *μ*_1_ is a set of controls that stabilizes *F*_1_ in 𝒞_1_ and *μ*_2_ is a control that stabilizes 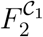 in 𝒞_2_, then *μ* = *μ*_1_ ∪ *μ*_2_ is a set of control that stabilizes *F* in 𝒞 = 𝒞_1_ ⊕ 𝒞_2_, as long as 𝒞_1_ or 𝒞_2_ is a steady state.

A recently published multicellular Boolean network model describes the microenvironment of pancreatic cancer cells by modeling the interactions of pancreatic cancer cells (PCCs), pancreatic stellate cells (PSCs), and their connecting cytokines (*30*). This network has 69 nodes, 114 edges, and possesses three non-trivial modules (Fig. 5a). Fig. 5b shows the directed acyclic graph, which describes the connections between the modules.

**Figure 5:**
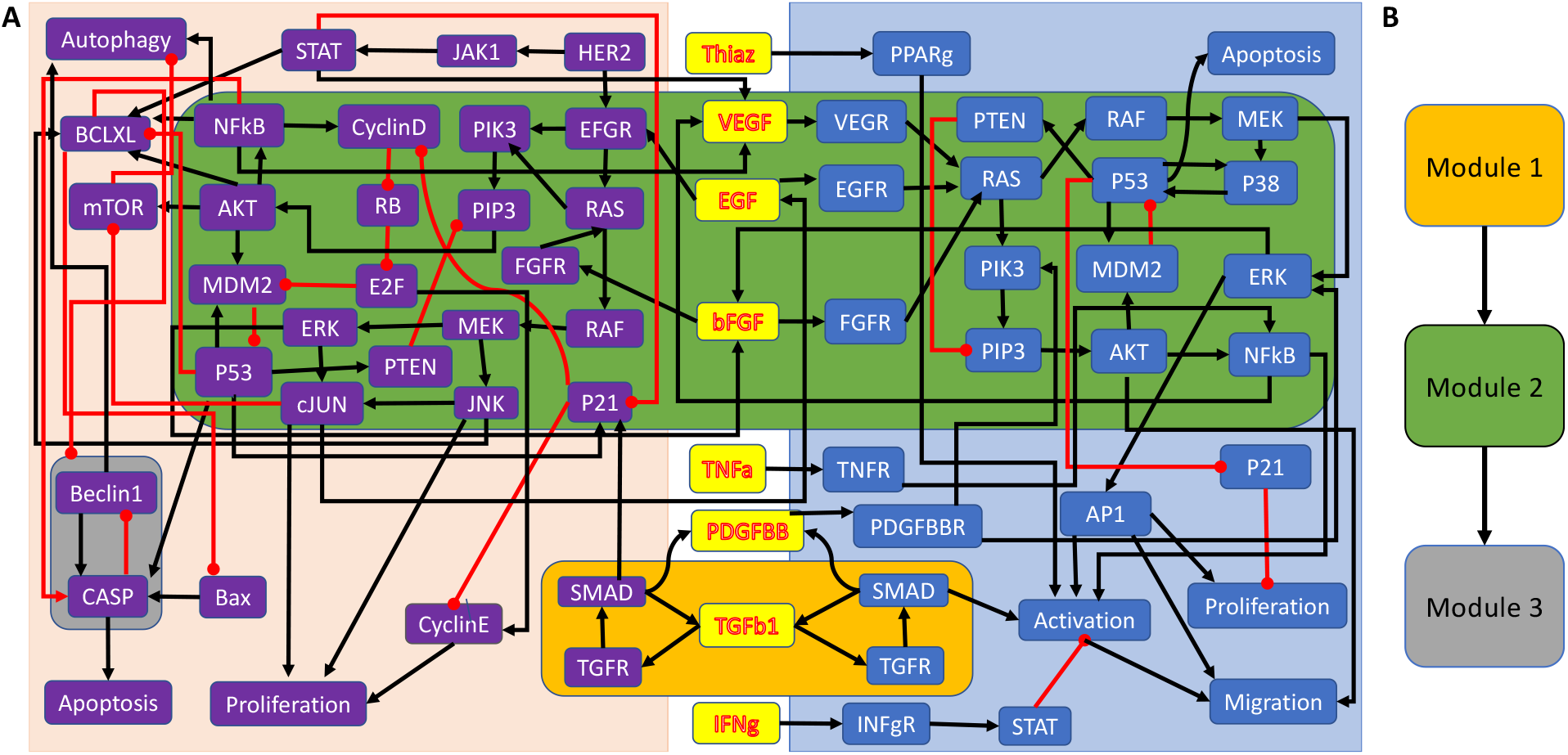
A multicellular Boolean cancer model (*30*), which describes the interactions of pancreatic cancer cells (purple nodes), pancreatic stellate cells (blue nodes), and their connecting cytokines (yellow nodes). (a) Wiring diagram describing the regulations between nodes, which are all monotonic, with black and red arrows indicating activation and inhibition, respectively. The non-trivial modules are highlighted by amber, green, and gray boxes. (b) Directed acyclic graph describing the connections between the non-trivial modules.

An effective treatment should induce the cancer cell to undergo apoptosis, which therefore represents the desired attractor of this network. To find a set of controls that stabilizes the network in this attractor, one can exploit the structural decomposition of the network by first controlling the upstream module (module 1), which has four attractors: two steady states and two 3-cycles. This module consists of two feedback loops joined by the node *TGFb*1. It is thus enough to control *TGFb*1 to stabilize this module into any of its attractors (*31*). Using the methods from (*32*) or (*26*), the controls of module 2 can be identified. A minimal set of two nodes needs to be controlled to stabilize this module: *RAS* in the pancreatic cell and *RAS* in the stellate cell. After applying these controls, the nodes in the downstream module (module 3) are all already constant and do therefore not require additional controls. Using the modular structure of the network, three nodes can be easily identified, which suffice to control the entire network. Notably, this never requires the consideration of the entire network, which saves computation time. Disregarding the decomposition and identifying controls for the whole network instead yields the same minimal set of three controls. However, this may not always be the case. In rare cases, the module-by-module control identification strategy will yield a set of controls that is larger than necessary.

## Discussion

The search for “fundamental laws” has been part of systems biology since its beginning, including features of biological systems that are characteristic of most or all systems of a given type, such as gene regulatory networks. The concept of modularity can be considered as such a feature, and has been studied extensively in several different contexts. Another focus of interest has been the relationship between the structure and function of dynamic networks. The results in this paper in essence provide evidence that modularity is in fact a key feature that connects structure and function of networks.

Systems biology has been a field that is making extensive use of mathematical models as descriptive language and analytic tool. Notions such as dynamic modularity are difficult or impossible to study without the use of mathematical models, as is the relationship between structure and function of networks. A limitation of this approach is of course that published models are partial and simplified representations of the requisite biology, so that caution is required when drawing conclusions. But this approach has yielded useful results in studying motifs in static networks (e.g., (*20*)). The advantage of a mathematical foundation is that it enables an analytical treatment of concepts that might otherwise have to be studied using heuristics, examples, and simulations. This is the essence of our approach in this study. Based on rigorous definitions, we were able to prove the link between structural and functional modularity, as well as the broad application to control of networks. We believe that we have only scratched the surface of results that follow from the mathematical framework we have established. For instance, the flip side of network decomposition is network construction through “concatenation” of modules. This can be done in ways that achieve certain dynamic properties, of potential interest to problems in synthetic biology.

Finally, while we have provided evidence that our concept of structural and functional modularity might have biological relevance, more work remains to be done. For instance, it would be of interest to investigate the biological features of the individual modules found in the repository of Boolean network models from (*14*) to investigate whether modules in our definition can be viewed as meaningful biological “functional units.” The implications of a functional modular structure also remain to be explored beyond our initial result of control at the modular level. We also believe that many of our results should hold in appropriate form for the modeling framework of ordinary differential equations.

## Methods

### Meta-analysis of published gene regulatory network models

We used the same repository of 122 published and distinct gene regulatory network models as in (*14*). Some of these models include non-essential regulators. That is, a node is included as a regulator in an update rule but a change in this node never affects the update rule. We removed all non-essential regulators from the update rules, before computing for each network the number of genes (i.e., size), the average connectivity, all SCCs, as well as the size of each SCC. From this, we derived the primary metric of interest, the number of non-trivial SCCs. Trivial SCCs consist of one node only. Since SCCs are defined as the largest connected component such that there is a path from every node to every *other* node, it is irrelevant whether the single node in a trivial SCC regulates itself.

The logistic multivariable regression model, implemented in the Python module statsmodels.api is given by

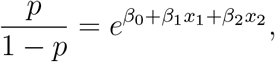

where *p* is the probability of a model having multiple non-trivial SCCs, and *x*_1_, *x*_2_ are average connectivity and network size.

### Generation of Boolean networks for simulation study

To understand the effect of modularity on the phenotypical robustness and the dynamical complexity, we resorted to simulation studies of Boolean networks with a specific structure and a defined number of SCCs (i.e., modules). To reduce the number of potential confounders, we fixed the network size at *N* = 60 and the in-degree of each node at *n* = 3, which is slightly higher than the average in-degree in published gene regulatory network models (*14*). We further considered only nested canalizing update rules since most rules in published gene regulatory networks are of this type (*14*). To generate networks with a defined number of *m* ∈ {1, 2, …, 6} modules, each of which consists of *N/m* nodes, we first generated a random directed acyclic graph of *m* modules by picking uniformly at random a weakly-connected lower triangular binary *m × m*-matrix *D* with diagonal entries 1. If *D*_*ij*_ = 1, a node in module *i* regulates a node in module *j*. Otherwise, there is no connection. To ensure that the number of SCCs was indeed *m*, we required each module to be a single SCC. We achieved this by randomly generating wiring diagrams for a module until the wiring diagram was strongly connected (for the sparsest modules (i.e., *m* = 1, *N/m* = 60), this took on average about 22 iterations).

### Estimating dynamical complexity and phenotypical robustness

The size of the state space of the 60-node Boolean networks used in the computational study prohibits the exhaustive identification of all attractors. To compute all attractors, we could have exploited the decomposition into smaller modules for decomposable networks. However, this does not help with the identification of attractors for non-decomposable networks consisting of a single module of size 60. To avoid introducing any bias by using different methods, we employed the same sampling technique to estimate a lower bound of the number of attractors for each Boolean network. Specifically for each network *F*, we generated 500 random initial states **x**_**0**_ ∈ {0, 1}^60^ and continued to synchronously update each **x**_0_ according to *F* (that is, **x**_*i*+1_ = *F* (**x**_*i*_)) until a recurring state was found, indicating the arrival at an attractor.

Biologically meaningful attractors “attract” a substantial portion of the state space. With a state space size of 2^60^ and when starting from 500 random initial states, we have a 95% chance of finding an attractor, which attracts 0.6% of the state space and even a 99% chance of finding an attractor, which attracts 0.9% of the state space. Relying on sampling and the resulting lower bound of the number of attractors should therefore not limit the validity of our findings.

To estimate the phenotypical robustness, we considered the same 500 random initial states **x**_0_ ∈ {0, 1}^60^ and generated for each **x**_0_ a corresponding state *y*_0_ = **x**_0_ ⊕*e*_*i*_ by randomly flipping one bit *i* ∈ {1, …, *n*} (where *e*_*i*_ is the *i*th unit vector and ⊕ denotes binary addition). Just as **x**_0_, we synchronously updated **y**_0_ according to *F* until it reached an attractor and compared the attractors. As a consequence, all estimated phenotypical robustness values are multiples of 1*/*500.

## Acknowledgments

The authors thank Elena Dimitrova for participating in initial fruitful discussions.

## Author contributions

C.K., M.W., A.V., D.M., and R.L. designed research, performed research, and wrote the paper; C.K., M.W., and D.M. analyzed data.

## Data availability

All data needed to evaluate the conclusions in the paper are present in the paper and/or the Supplementary Materials.

## Supplementary Material

### 1 Proofs of the structural decomposition theorems

This section contains the proofs of the structural decomposition theorems described in the main text. First, we define in full detail the semi-direct product, used to combine two networks in a hierarchical fashion.

#### Definition 1.1.

Consider two Boolean networks,

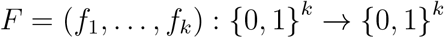

with variables *x* = (*x*_1_, …, *x*_*k*_) and

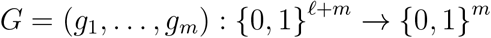

with external inputs *u* = (*u*_1_, …, *u*_*ℓ*_) and variables *y* = (*y*_1_, …, *y*_*m*_). Let Λ ⊆ {1, …, *k*} such that |Λ| = *f*, and define 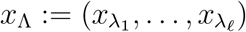. Then,

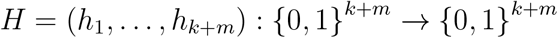

defines a combined Boolean network by setting

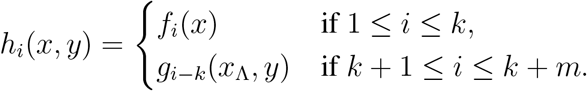

That is, the variables *x*_Λ_ act as the external inputs of *G*. The corresponding coupling scheme is defined to be

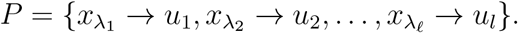

We denote *H* as *H* := *F* ⋊_*P*_ *G* and refer to this as the coupling of *F* and *G* by (the coupling scheme) *P* or as the semi-direct product of *F* and *G* via *P* .

#### Theorem 1.1.

If a Boolean network *F* is not a module, then there exist *F*_1_, *F*_2_, *P* such that

*F* = *F*_1_ ⋊_*P*_ *F*_2_. Furthermore, we can find a decomposition such that *F*_1_ is a module.

*Proof*. Let *F* = (*f*_1_, …, *f*_*n*_) be a Boolean network with variables *X* = {*x*_1_, …, *x*_*n*_} and assume *F* is not a module. Then the wiring diagram of *F* is not strongly connected, implying there exists at least one node *y* and one node *x*_*j*_ ≠ *y* such that there exists no path from *x*_*j*_ to *y* in the wiring diagram of *F* . Let 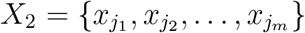 denote the set of all such nodes, i.e., the nodes for which there exists no paths to *y*. Further, let *X*_1_ = *X\X*_2_ denote the complement set of nodes to *X*_2_. Note that for every *x*_*i*_ ∈ *X*_1_, there exists a path from *x*_*i*_ to *y* but no paths originating from *X*_2_ to *x*_*i*_.

Define Λ to be the subset of indices Λ = {*λ*_1_, …, *λ*_*ℓ*_} ⊂ {1, …, *k*} such that for each *λ* ∈ Λ there exists at least one function 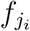 with 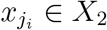 which depends on *x*_*λ*_.

If Λ = ∅, then the sets *X*_1_ and *X*_2_ represent two groups of nodes, which are disconnected in the wiring diagram. Hence the network *F* is a Cartesian product of *F*_1_ and *F*_2_. It follows that *F* = *F*_1_ ⋊_*P*_ *F*_2_ with *P* = ∅.

If Λ ≠ ∅, then for any *x*_*i*_ ∈ *X*_1_, the corresponding update function *f*_*i*_ does not depend on *X*_2_ by construction, as there are no paths from *X*_2_ to *x*_*i*_, and we set *F*_1_ to be the restriction of *F* to 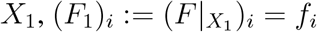. For any *x*_*i*_ ∈ *X*_2_, if the corresponding update function depends on a node *x*_*j*_ ∈ *X*_1_, then *x*_*j*_ ∈ Λ by the definition of Λ. It follows by construction that any function *f*_*i*_ then can be written as a Boolean function on *X*_2_ with external inputs from *x*_Λ_.

Hence, *F* = *F*_1_ ⋊_*P*_ *F*_2_.

Note that in the above proof we can choose the node *y* such that it belongs to a SCC that receives no edge from any other SCC. *X*_1_ will contain the nodes of this SCC and hence *F*_1_ will be a module. □

The main structural decomposition theorem follows directly from this:

#### Theorem 1.2.

For any Boolean network *F*, there exist unique modules *F*_1_, …, *F*_*m*_ such that

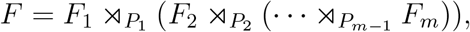

where this representation is unique up to a reordering, which respects the partial order of *Q* (Eq. 1), and the collection of coupling schemes *P*_1_, …, *P*_*m*−1_ depends on the particular choice of ordering.

*Proof*. If *F* is a module, then *m* = 1 and the result follows.

If *F* is not a module, we use induction on the downstream subnetwork *F*_2_ in Theorem 1.1 to obtain the result. □

### 2 Non-autonomous Boolean networks

This section contains the full definition of non-autonomous Boolean networks, as well as two examples.

#### Definition 2.1.

A non-autonomous Boolean network is defined by

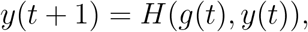

where *H* : {0, 1}^*k*+*m*^ → {0, 1}^*m*^ and 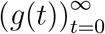 is a sequence with elements in {0, 1}^*k*^. The network, denoted *H*^*g*^, is non-autonomous because its dynamics depend on *g*(*t*).

A state *c* ∈ {0, 1}^*n*^ is a *steady state* of *H*^*g*^ if *H*(*g*(*t*), *c*) = *c* for all *t*. Similarly, an ordered set with *r* elements, *𝒞* = {*c*_1_, …, *c*_*r*_} is an *attractor of length r* of *H*^*g*^ if *c*_2_ = *H*(*g*(1), *c*_1_), *c*_3_ = *H*(*g*(2), *c*_2_), …, *c*_*r*_ = *H*(*g*(*r* − 1), *c*_*r*−1_), *c*_1_ = *H*(*g*(*r*), *c*_*r*_), *c*_2_ = *H*(*g*(*r* + 1), *c*_1_), In general, *g*(*t*) is not necessarily of period *r* and may even not be periodic.

If *H*(*g*(*t*), *y*) = *G*(*y*) for some network *G* for all *t* (that is, it does not depend on *g*(*t*)), then *y*(*t* + 1) = *H*(*g*(*t*), *y*(*t*)) = *G*(*y*(*t*)) and this definition of attractors coincides with the classical definition of attractors for (autonomous) Boolean networks.

#### Example 2.1.

Consider the non-autonomous network defined by

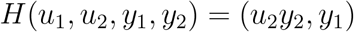

and the two-periodic sequence 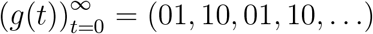, which corresponds to a 2-cycle of the upstream 2-node network. If the initial point is 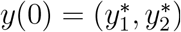, then the dynamics of *H*^*g*^ can be computed as follows:

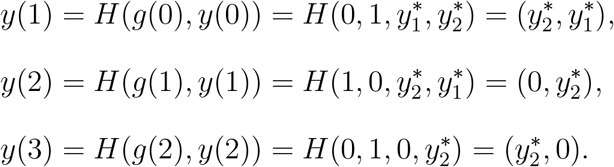

Thus for 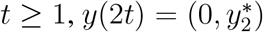 and 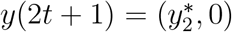. It follows that the attractors of *H*^*g*^ are given by 00 (one steady state) and (01, 10) (one cycle of length 2). Note that (10, 01) is not an attractor because (10, 01, 10, 01, …) is not a trajectory for this non-autonomous network. This is a subtle situation that can be sometimes missed when not considering all trajectories a limit cycle represents.

#### Example 2.2.

Consider the non-autonomous network defined by *H*(*u*_1_, *u*_2_, *y*_1_, *y*_2_) = (*u*_2_*y*_2_, *y*_1_), as in the previous example, and the one-periodic sequence 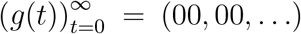, which corresponds to a steady state of the upstream 2-node network. If the initial point is 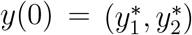, then the dynamics of *H*^*g*^ can be computed as follows:

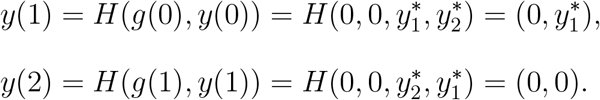

Then, *y*(*t*) = (0, 0) for *t* ≥ 2, and the only attractor of *H*^*g*^ is the steady state 00.

### 3 Proof of the dynamic decomposition theorem

For a decomposable network *F* = *F*_1_ ⋊_*P*_ *F*_2_, we introduce the following notation for attractors. First, note that *F* has the form *F* (*x, y*) = (*F*_1_(*x*), *F*_2_(*x, y*)) where *F*_2_ is a non-autonomous network. Let 𝒞_1_ = (*r*_1_, …, *r*_*m*_) ∈ 𝒜 (*F*_1_) and 𝒞_2_ = (*s*_1_, …, *s*_*n*_) ∈ 𝒜 (*F*_1_) ∈ 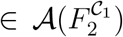 be attractors of length *m* and *n*, respectively. Then, the sequence 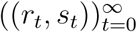 has period *l* = *lcm*(*m, n*), so we define the sum (or concatenation) of these attractors to be

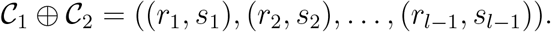

Note that the sum of attractors is not a Cartesian product, 𝒞_1_ *×* 𝒞_2_ = {(*r*_*i*_, *s*_*j*_)| for all *i, j*}.

Similarly, for an attractor 𝒞_1_ and a collection of attractors *A* we define

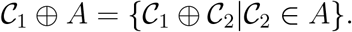

Our second main theoretical result shows that the dynamics (i.e., the attractor space) of a semi-direct product can be seen as a type of semi-direct product of the dynamics of the decomposable subnetworks. When applied iteratively, this enables a computation of the attractor space from the attractor space of each module.

#### Theorem 3.1.

Let *F* = *F*_1_ ⋊_*P*_ *F*_2_ be a decomposable network. Then

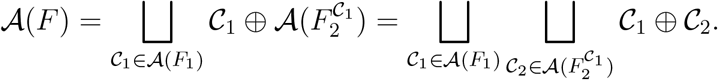

*Proof*. Let *X*_1_ and *X*_2_ be the variables of *F*_1_ and *F*_2_, respectively. Further, let 𝒞 = {*c*_1_, …, *c*_*l*_} ∈ 𝒜 (*F*) be an arbitrary attractor of *F* with length *l*. We can define 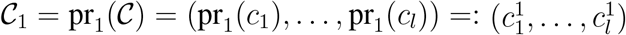 as the projection of 𝒞 onto *X*_1_, and similarly 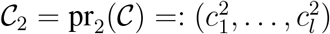 as the projection of 𝒞 onto *X*_2_. By definition, *F*_1_ does not depend on *X*_2_. Thus, *F*_1_(pr_1_(*x*)) = pr_1_(*F* (*x*)), and for any 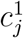,

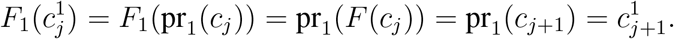

Iterating this, we find that in general 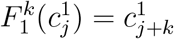, from which it follows that 𝒞_1_ ∈ 𝒜 (*F*_1_).

Next, we consider the non-autonomous network 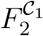 defined as in Definition 2.1 where *y*(*t* + 1) = pr_2_*F* (*g*(*t*), *y*(*t*)), and 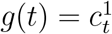. If 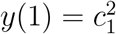, then

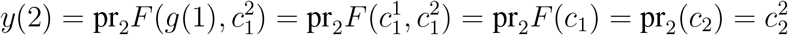

and in general

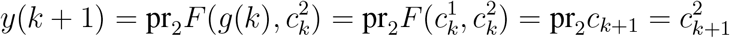

Hence *y*(*l* + 1) = pr_2_*F* (*c*_*l*_) = pr_2_*c*_1_ = *y*(1) and thus 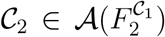. From this we have that 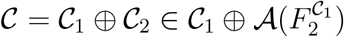 and thus

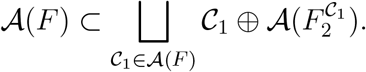

Conversely, let 𝒞_1_ ∈ 𝒜 (*F*_1_) and 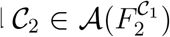. We want to show that 𝒞_1_ ⊕ 𝒞_2_ ∈ 𝒜 (*F*). Let 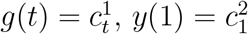, and *y*(*t* + 1) = pr_2_*F* (*g*(*t*), *y*(*t*)). Since 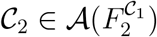, then 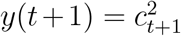 by definition. Let *N* = |𝒞_2_|. Then

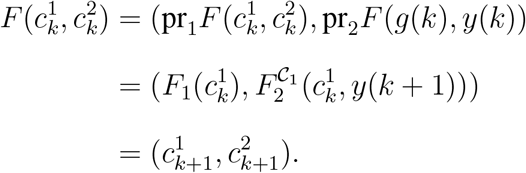

Thus 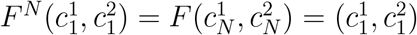 and hence 𝒞_1_ ⊕ 𝒞_2_ ∈ 𝒜 (*F*). It follows that

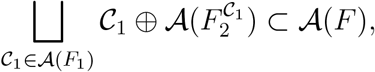

from which we can conclude that the sets are equal. □

The following two examples highlight how Theorem 3.1 enables the computation of the dynamics of a decomposable network from the dynamics of its modules. To match attractors from the upstream module with the attractor spaces of the corresponding non-autonomous downstream networks, it is useful to consider the space of attractors in a specified order: we use parentheses (curly brackets) to denote an ordered (unordered) space of attractors. If there is no ambiguity, in practice we can use ⋊ instead of ⋊_*P*_.

#### Example 3.1.

Consider the Boolean network *F* (*x*_1_, *x*_2_, *y*_1_, *y*_2_) = (*x*_2_, *x*_1_, *x*_2_*y*_2_, *y*_1_). We can decompose *F* = *F*_1_ ⋊ *F*_2_ where *F*_1_(*x*_1_, *x*_2_) = (*x*_2_, *x*_1_) is an upstream module and *F*_2_(*u*_2_, *y*_1_, *y*_2_) = (*u*_2_*y*_2_, *y*_1_) is a downstream module with external parameter *x*_2_. To find all attractors of *F* by using Theorem 3.1, we find the attractors of *F*_1_ and the attractors of *F*_2_ induced by each of those attractors. It is easy to see that 𝒜 (*F*_1_) = {00, 11, {01, 10}} (where we denote steady states 𝒞 = {*c*} simply by *c*).

- For 𝒞_1_ = 00, the corresponding non-autonomous network is *y*(*t* + 1) = *F*_2_(0, 0, *y*(*t*)). If 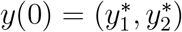, then

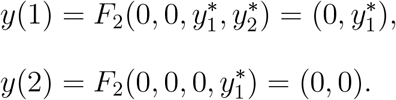

Thus, the space of attractors for 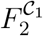 is

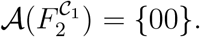
- For 𝒞_2_ = 11, the corresponding non-autonomous network is *y*(*t* + 1) = *F*_2_(1, 1, *y*(*t*)). If 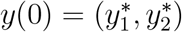, then

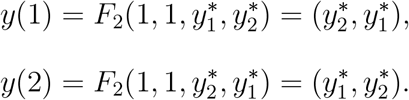

Thus, the corresponding space of attractor is

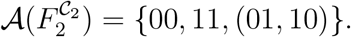
- For 𝒞_3_ = (01, 10), we define *g*(*t*) : ℕ→ {0, 1}^2^ by *g*(0) = (0, 1), *g*(1) = (1, 0), and 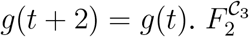 is given by *y*(*t* + 1) = *F*_2_(*g*(*t*), *y*(*t*)). If 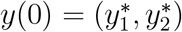, then

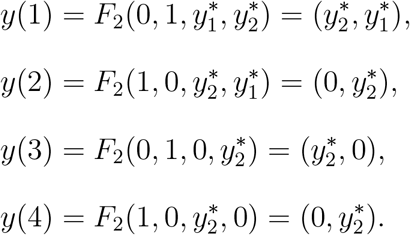

Then, the corresponding space of attractors is

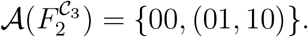

To reconstruct the entire space of attractors for *F*, we have

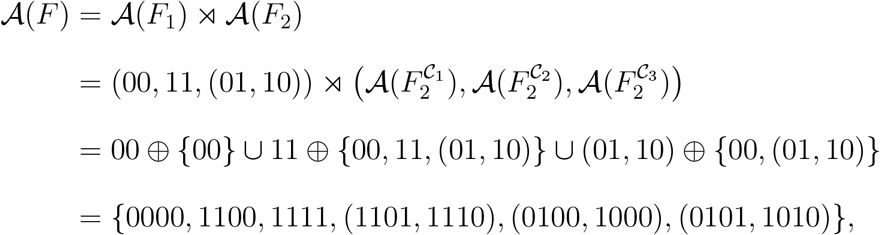

which agrees with the space of attractors shown in Fig. 4B.

#### Example 3.2.

Consider the linear Boolean network

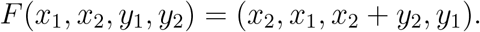

We can decompose *F* = *F*_1_ ⋊ *F*_2_ into modules *F*_1_(*x*_1_, *x*_2_) = (*x*_2_, *x*_1_) and *F*_2_(*u*_2_, *y*_1_, *y*_2_) = (*u*_2_ + *y*_2_, *y*_1_). The space of attractors of the upstream module *F*_1_ is

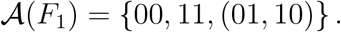

Using the dynamic decomposition theorem (Theorem 3.1), we can identify all attractors of *F* as follows (see Fig. S2 for a graphical description).

- For 𝒞_1_ = 00, the corresponding non-autonomous network is 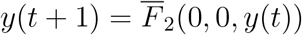. If 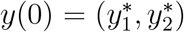, then 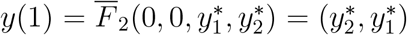. Thus, the space of attractors for 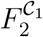 is

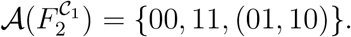
- Similarly, for 𝒞_2_ = 11, we find that the space of attractors for 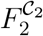 is

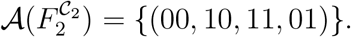
- For 𝒞_3_ = (01, 10), we define *g*(*t*) : ℕ→ *X*_1_ by *g*(0) = (0, 1), *g*(1) = (1, 0), and 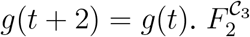 is given by *y*(*t* + 1) = *F*_2_(*g*(*t*), *y*(*t*)). If 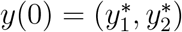, then

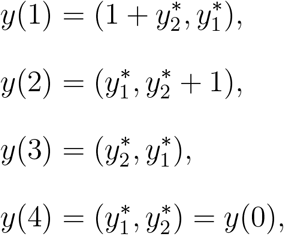

and in general for *t >* 0,

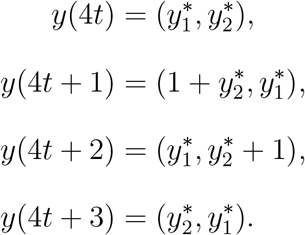

It follows that there are only 2 periodic trajectories in this case: (00, 10, 01, 00, 00, 10, 01, 00, …) and (11, 01, 10, 11, 11, 01, 10, 11, …), which both have period 4. The corresponding attractor space is

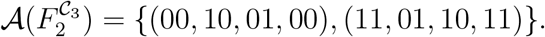

Note that the repetition of certain states is needed to obtain the correct attractors of the full network *F* .

To reconstruct the space of all attractors for *F*, we have

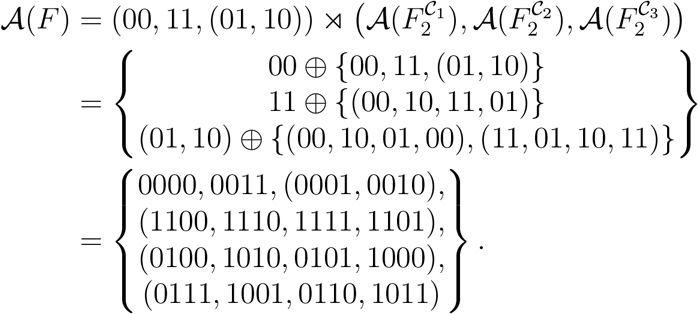

The linear network *F* possesses thus two steady states, one 2-cycle and three 4-cycles.

**Figure S1:**
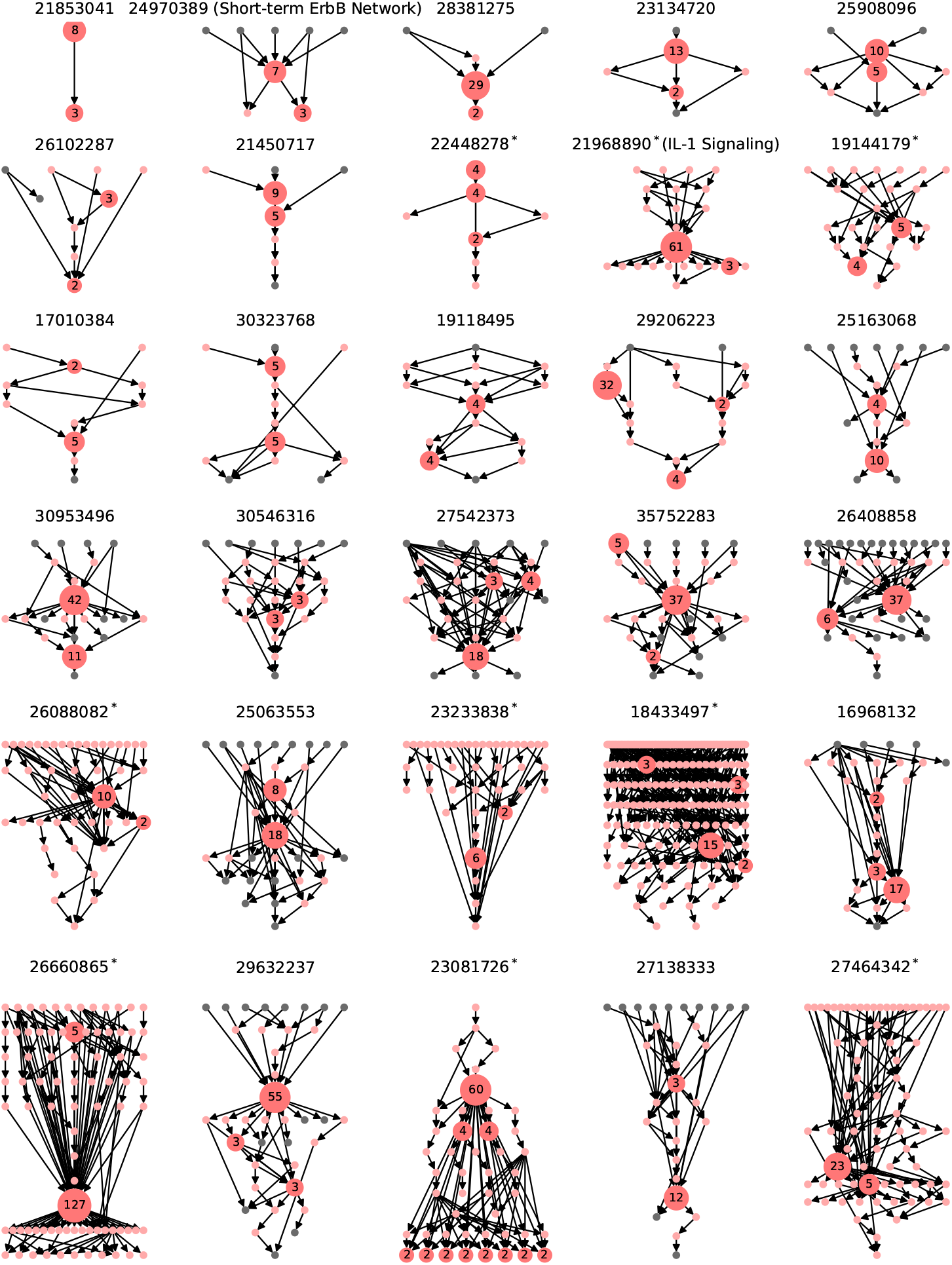
Modular decomposition of all published expert-curated Boolean gene regulatory network models with more than one non-trivial module. Each model is labeled by the Pubmed ID of its source. Each red non-trivial module is labeled by its size, i.e., the number of nodes contained in the module. Trivial modules consist of one node only. They are colored gray if they are input or output nodes, i.e., nodes without incoming or outgoing edges, respectively. Otherwise, they are colored pink. For models with more than 40 modules, input and output modules are omitted for clarity, indicated by ^∗^ after the Pubmed ID. An arrow from module *X* to module *Y* indicates that some node in *X* regulates some node in *Y* . The directed acyclic graph of the multicellular pancreatic cancer model, analyzed in Fig. 5, is shown in row 4, column 4 (Pubmed ID 35752283).

**Figure S2:**
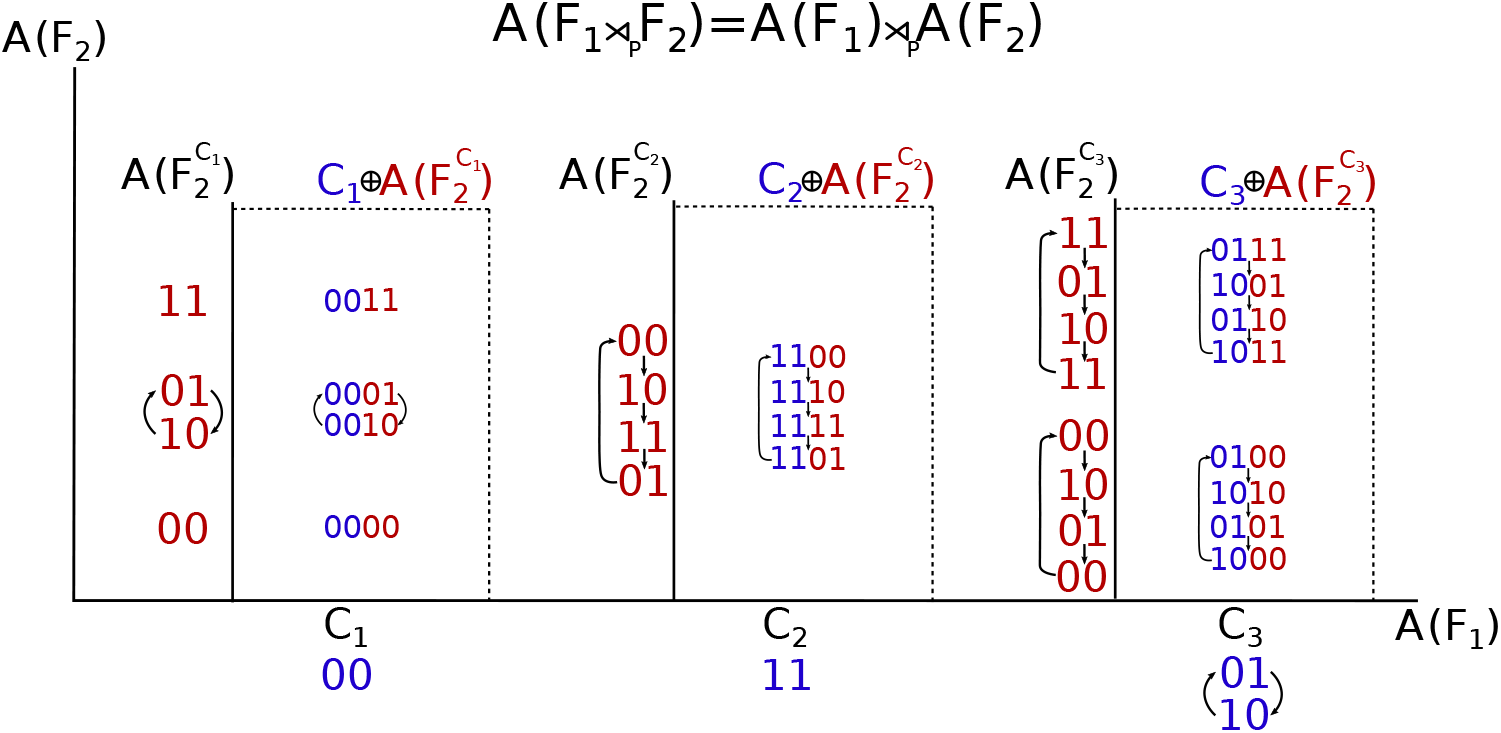
Graphical description of the dynamic decomposition theorem (applied to Example 3.2). The dynamics of *F*_1_ ⋊_*P*_ *F*_2_ can be seen as a semi-direct product between the dynamics of *F*_1_ and the dynamics of *F*_2_ induced by *F*_1_ via the coupling scheme *P* . The dynamics of *F*_2_ induced by attractors of *F*_1_ can vary, and the dynamic decomposition theorem (Theorem 3.1) shows precisely how to combine all of these attractors.

